# Measuring the similarity of SMLM-derived point-clouds

**DOI:** 10.1101/2022.09.12.507560

**Authors:** Mohammed Baragilly, Daniel J. Nieves, David J. Williamson, Ruby Peters, Dylan M. Owen

## Abstract

Single-molecule localisation microscopy produces data in the form of point-clouds. Here, we present a tool for assessing the similarity of two such point-clouds, which, unlike measures such as co-localisation, is insensitive to differences that are not preserved between data sets. The presented method can determine whether two point-clouds were generated from the same conditions and can identify from which of two experimental conditions an unseen point-cloud was likely derived.

## Introduction

Single-molecule localisation microscopy (SMLM) outputs data in the form of a point-cloud representing^1^. These can either be rendered to produce images or interrogated directly to extract biologically relevant information. For the latter, various methods of cluster or fibre analysis have been developed. An important capability is to be able to quantify the similarity two point-clouds for which there are several methods, depending on the measure required. For example, a number of co-localisation and co-clustering measures have been developed, notably the Cross-Ripley’s K-function and Pair-Correlation function^2–5^. These measures require the two point-clouds to be derived from the *same space* to be meaningful – for example, co-localisation is measured between two channels acquired in the same region-of-interest (ROI) within the same cell. A separate class of similarity measures exist when the two point-clouds come from different cells or conditions. The most common measure is to test whether statistical descriptors of the point-clouds are similar. For example, users may perform cluster analysis and compare the sizes of clusters, the number of clusters etc^6^,^7^. This approach does not produce a similarity measure as there is no defined way to combine these descriptive outputs into one overall score.

We present a non-parametric statistical approach to assess similarity between two point-clouds derived from different ROIs that provides a score independent of specific statistical descriptors. Since it is designed for ROIs collected separately (e.g. from different cells), it is insensitive to differences that are not preserved in the context of cell biology or SMLM, such as the positions of clusters within the point cloud and rotations of the ROI. We show that being able to extract a single dissimilarity score from otherwise complex data, is profoundly useful. For example: the method can take two sets of simulated point-clouds and determine if they were generated using different simulation conditions. The biological analogy to this, which we also demonstrate, is the assessment of similarity between two point-clouds derived from two experimental conditions, for example, a wild-type condition versus a mutant. This approach generates a p-value, reporting on the significance of similarity, i.e. whether a treatment has an effect. The method can also take a single ROI and determine from which of a selection of conditions it was most likely derived. Together, our method provides a simple means of assessing similarly in complex point-cloud data, in a quantitative, robust and rapid manner.

## Results

We have developed a similarity measure for comparing two point-clouds, described mathematically in the Online Methods. Briefly, we divide a ROI into 30 nm pixels and calculate the cumulative frequency histogram for the number of localisations per pixel. A Kolmogorov-Smirnov (K-S) test is then used to derive a similarity measure between two such histograms in the form of a dissimilarity value, which we term λ (Figure 1b). A value of zero indicates that the point-clouds are identical. Higher values indicate increasing dissimilarity. Typically, multiple ROIs will be acquired per condition. In this case, we compute all possible pair-wise λ-values both within and between conditions, from which we derive a histogram of λ-values, and the mean, 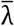. This allows for a p-value calculation (Figure 1b).

**Figure 1:**
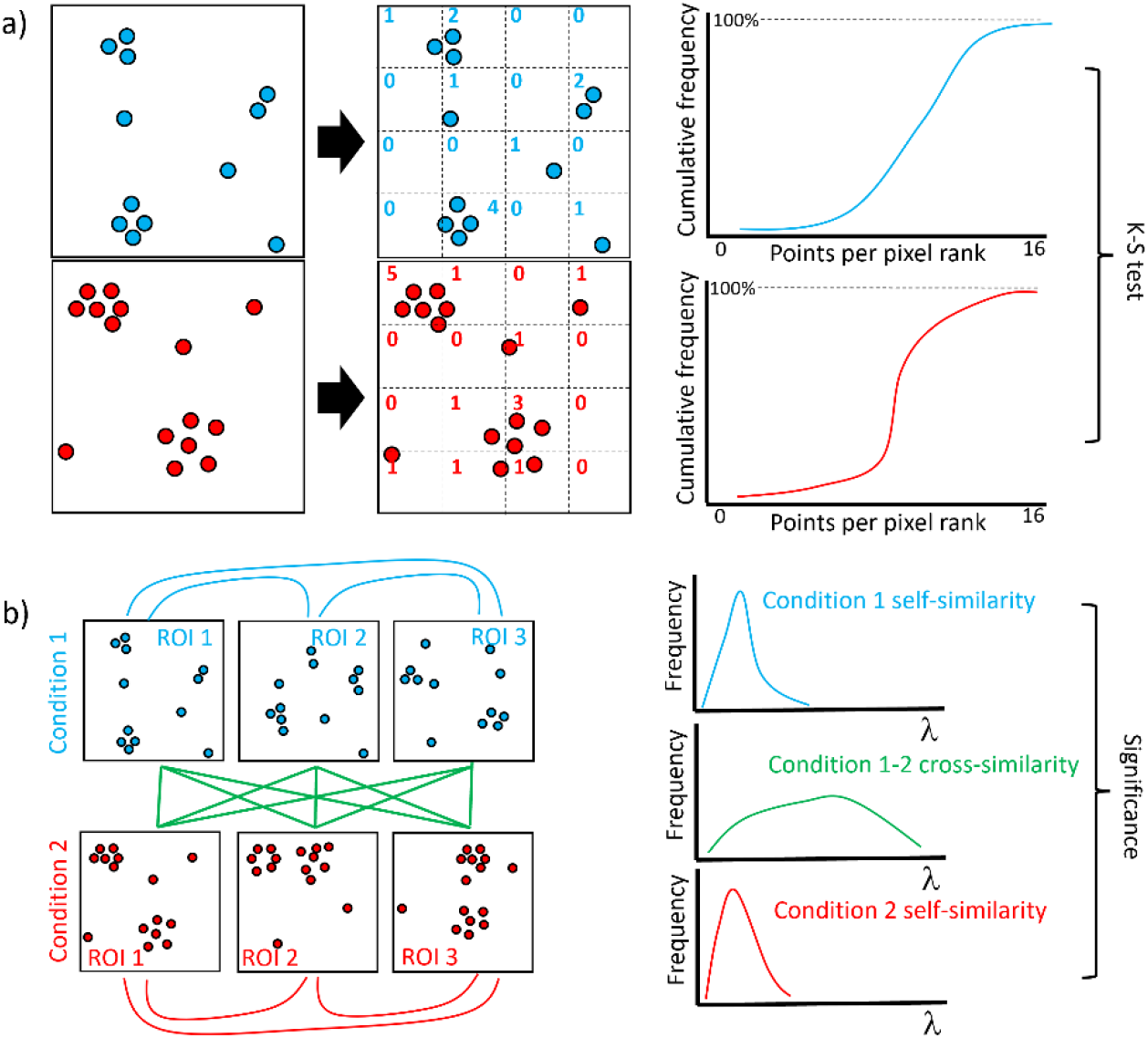
Description of the algorithm. a) Schematic of the algorithm which relies on a K-S test between the cumulative frequency histograms of the number of localisations per 30 nm pixel. b) For multiple ROIs per condition, every pair-wise comparison is made, resulting in a histogram of λ-values to which significance testing can be applied.

We first demonstrate the method on simulated data. Localisations are placed into circular, Gaussian clusters whose centre positions are located randomly within an ROI. On top of these, un-clustered points are placed randomly within the ROI. Points are then scrambled to reflect the localisation error. A default condition generates 30 3000 x 3000 nm ROIs each containing 10 50nm clusters of 100 localisations and 1000 (50%) unclustered localisations, with an average localisation precision of 30 nm. A representative example is shown in Figure 2a. We first tested the algorithm by varying the cluster size and obtaining the dissimilarity scores with respect to the default condition (Figure 2b). We then varied the number of clusters per ROI (Figure 2c), the number of localisations per cluster (Figure 2d) and the percentage of localisations in the background (Figure 2e), and compared to the default condition. The algorithm was able to correctly determine that the given condition was significantly different from the default case. We also performed a comparison of point clouds derived from fibrous structures (Supplementary Figure 1) and showed that the algorithm can discriminate between different arrangements of fibres. Finally, the total number of points in the ROI may or may not be relevant and/or controlled in the experiment. We therefore developed a version of the code that is insensitive to the total number of localisations by randomly thinning the denser dataset (Supplementary Figure 2). They show that the algorithm can still distinguish between these conditions even when insensitive to overall point density. Note the results when varying the number of molecule per cluster give very low values because this simulation parameter only alters the overall point density and the algorithm therefore perceives these as similar, as expected.

**Figure 2:**
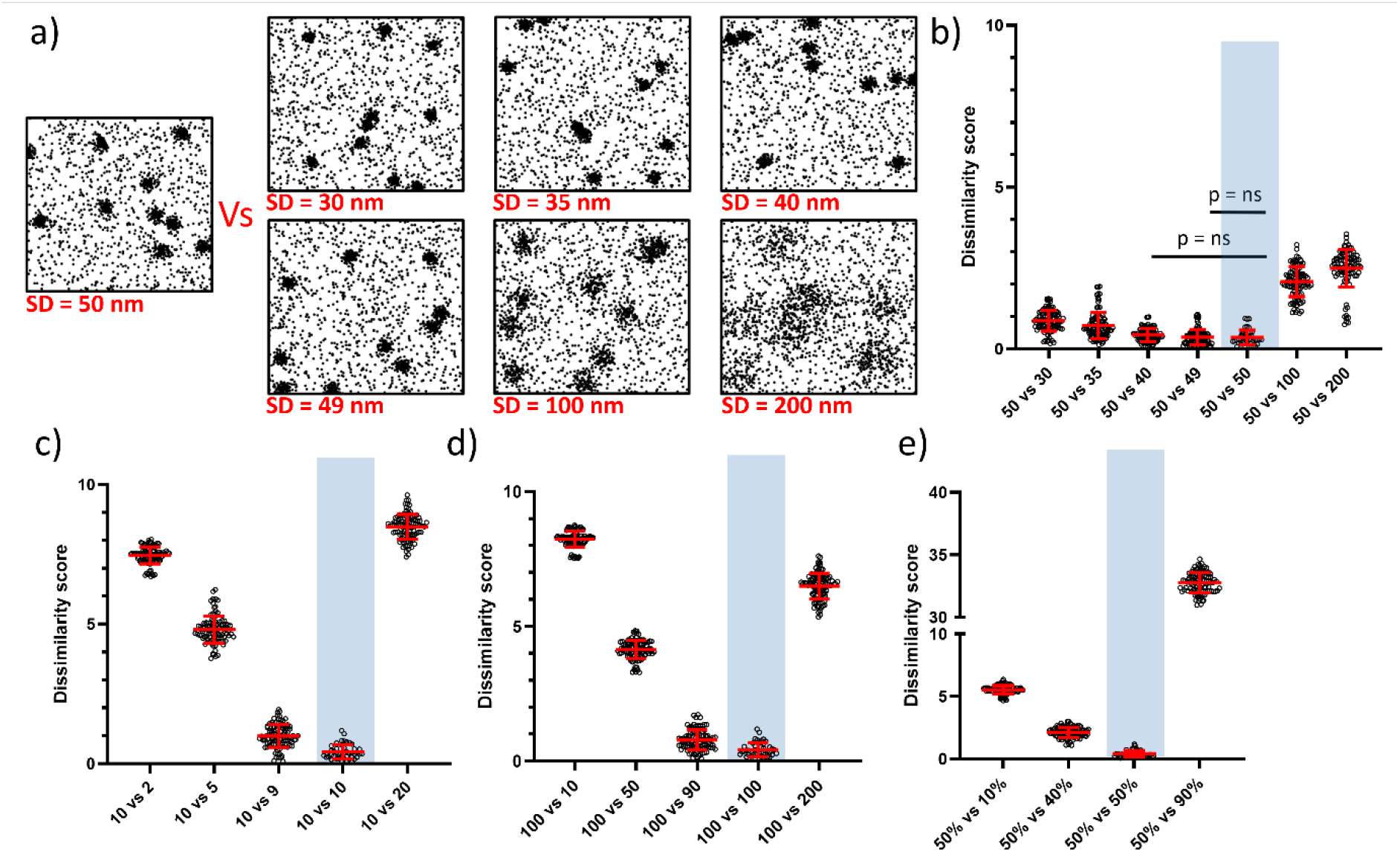
Demonstration of the method on simulated data. a) Representative simulated ROIs for the default condition and varying the cluster size (SD). Bee-swarm plots of λ-values for the default condition versus b) varying cluster sizes (nm). c) varying numbers of clusters per ROI. d) varying values of localisations per cluster. e) varying values for the percentage of un-clustered molecules. Bars show mean and S.D.

**Figure 2.**
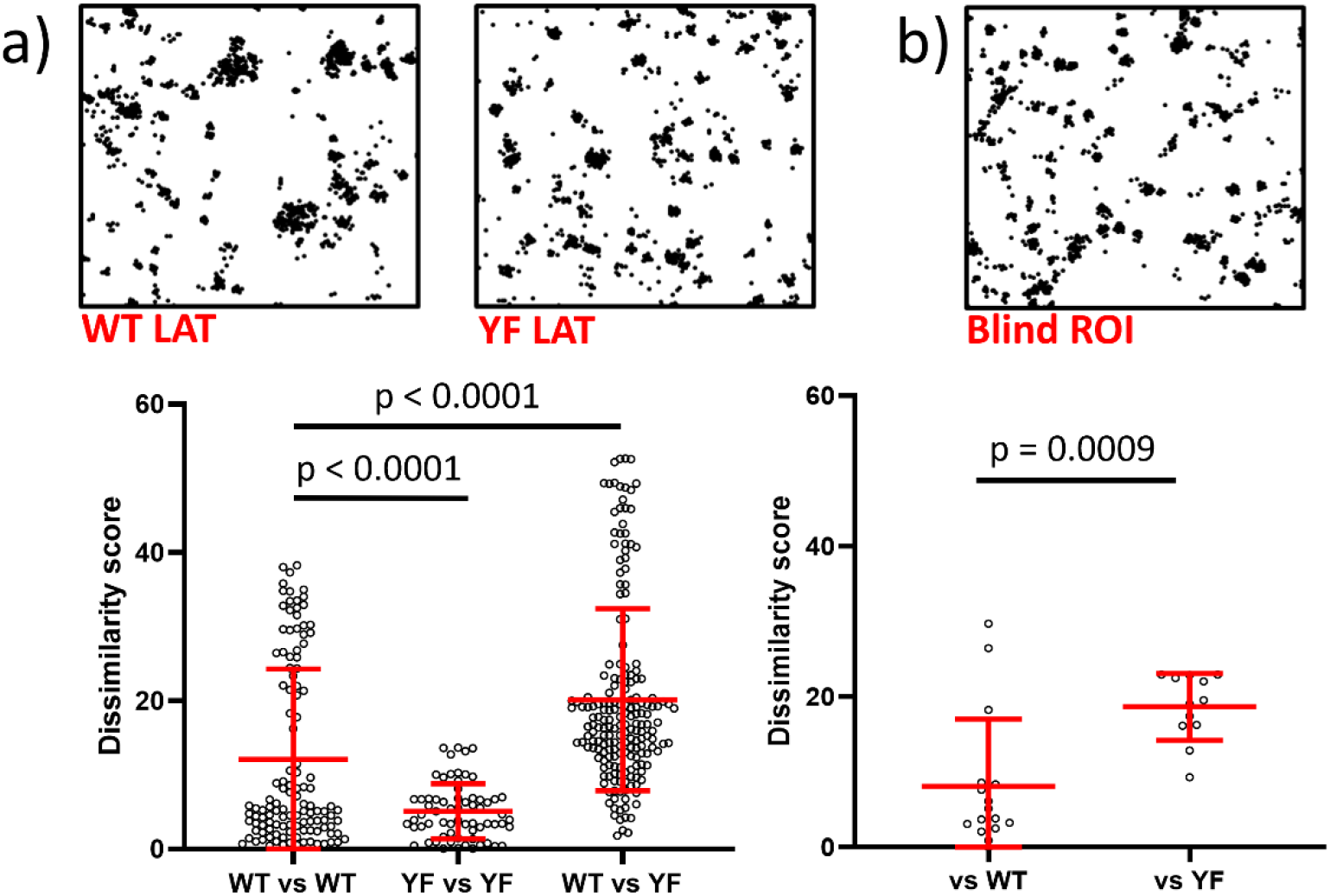
Demonstration of the method on biological data acquired via PALM. a) Representative ROIs from WT and YF LAT together with Bee-swarm plots for the dissimilarity scores within and between conditions. b) A blind ROI (example shown) is compared to all other ROIs in the WT and YF conditions. Bars show mean and S.D.

We next acquired PALM data of Linker-for-Activation of T cells (LAT)-mEos3.2 at artificial T cell synapses^8^. As a comparison, we generated a version of LAT lacking phosphorylatable tyrosine residues (YF LAT). λ-values for the WT-WT LAT comparison (Figure 3a) were significantly higher than for YF-YF LAT, indicating greater diversity of nanoscale organisation. λ-values for the WT-YF LAT comparison were significantly higher still, indicating that the YF LAT mutant does show differing nanoscale organisation. We next tested whether the algorithm, given a single blinded ROI, can determine from which condition it was most-likely derived. A blindly selected ROI is shown in Figure 3b, together with λ-values versus the other ROIs in the WT or YF condition.

Scores are significantly higher against the YF LAT condition, indicating the ROI likely derived from the WT condition; the correct result.

## Discussion

Obtaining a quantitative comparison of SMLM-derived point clouds is an important and necessary development to enable a direct, simple and robust means of comparing complex point-cloud data sets. Current methods exist on a spectrum – co-localisation measures, which compare point clouds derived from the same position and statistical comparison of extracted features between clouds from different conditions. Neither produces a similarity score between two separately acquired data sets. Here, we develop such a capability via an algorithm which is simple and rapid. The method allows the following new capabilities: a) to test whether an intervention (e.g. drug treatment, protein mutation) alters the nanoscale organisation of a given molecule. b) To assign a new ROI as have being derived from one of a number of previously acquired experimental conditions (e.g. from a healthy cell or a diseased cell). c) to detect anomalous data within a data collection, d) To tune the parameters of a data simulator to allow it to generate data statistically close to experimental data, but for which the ground truth distribution is known and e) to implement a search of a database of point-clouds for the most statistically similar condition.

## Supporting information

Software

## Acknowledgements

MB and DMO acknowledge funding from the Alan Turing Institute (ATI).

## Author contributions

MB developed the algorithm. MB analysed data. DJN, DJW and RP provided simulated and experimental data. MB and DMO conceived the work. MB and DMO wrote the manuscript. MB, DJN, DJW, RP and DMO edited the manuscript.

## Online methods

### Code availability

Code is available as Supplementary Material together with installation instructions.

### Sample preparation

Jurkat E6.1 cells (ECACC 88042803) expressing LAT-mEos3.2 were introduced to anti-CD3 (at 2 μg/ml; eBioscience clone OKT3, 16-0037-81) and anti-CD28 (at 5 μg/ml; RnD Systems, clone CD28.2, 16-0289-85) coated glass-bottomed chamber slides (#1.5 glass, ibidi μSlides) at 50 × 10^3^ cells/cm^2^ in warm HBSS and incubated at 37°C for 5 minutes to allow for synapse formation. Chamber wells were gently washed with warm HBSS and fixed in 3% paraformaldehyde for 20 minutes at 37°C. Fixed cells were washed in PBS.

### PALM imaging and image reconstruction

PALM image sequences were acquired on a Nikon N-STORM system in TIRF using a 100 × 1.49 NA CFI Apochromat TIRF objective running NIS Elements software v4.6. Samples were continuously illuminated with 561 nm laser light at approximately 2 kW/cm2 and 405 nm laser light at approximately 2 W/cm2. PALM data was acquired using an Andor IXON Ultra 897 EMCCD with an EM gain of 200 and pre-amplifier gain profile 3 to a centered 256 × 256 pixel region at 40 ms per frame for 5,000 to 15,000 frames. Localization of fluorophore coordinates were reconstructed using ThunderSTORM^9^ and corrected for sample drift using cross-correlation of images from five bins at a magnification of five.

### Description of the algorithm

Various statistical methods have been proposed in order to measure the similarity/dissimilarity between graphs. Some depend on finding the union and intersection for two or more graphs based on using the vertex labels. While this approach is straightforward, it can return different values if the vertices are relabelled without changing the structure of the graphs. The Jaccard index is one statistic used for measuring the similarity and diversity of sample sets based on union and intersection, but does not consider term frequency, instead, simply counting the number of terms that are common between two sets. Moreover, it ignores rare features in the structure of the data, which are significantly informative in the comparison process. Other similarity approaches have been used to classify nearest neighbours and measure the similarity between the sets. Examples of these approaches are cosine similarity, Pearson’s correlation, mean square distance, Neuman Kernels and several data mining algorithms. Many of these approaches are complex or computationally expensive. As a result, the graph edit distance (GED), which relies on structure rather than vertex labels, has been introduced to measure the similarity between pairwise graphs, error-tolerantly in inexact graph matching. However, computing the graph edit distance between two graphs is NP-complete problem. Finding the identity for each vertex and edge in two graphs is called the isomorphism problem. In order to overcome the limitations of GED, the A*-based (search) algorithm has been used as one of the most efficient algorithms for measuring the similarity between graphs and more specifically in network analysis. It estimates the cost of path-completion and finds the optimal with minimum cost. However, the A* algorithm suffers from complexity, which depends on the heuristic, especially in the unbounded search space case as the number of nodes expanded is exponential in the depth of the solution.

The Kolmogorov-Smirnov image comparison technique uses the image histogram and grayscale distribution to test whether two images are similar. Here, instead of calculating the grayscale distribution, we calculate the empirical cumulative distribution function for the number of points in each pixel. The proposed method is a data-driven, non-parametric distribution free method which is less sensitive to the statistical model assumptions. It is also straightforward and efficient in terms of the computational time.

Generally, let *X*_1_…, *X_m_* and *Y*_1_,…, *Y_n_* be two random samples from a continuous population, 1 and 2, with distribution functions F and G respectively, where both *X*’s and *Y*’s are mutually independent and identically distributed. Using a Kolmogorov-Smirnov test, our goal is to determine whether there are any differences between the *X* and *Y* probability distributions. In other words, we are interested in testing the null hypothesis:

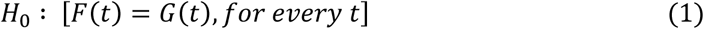

against the most general alternative hypothesis:

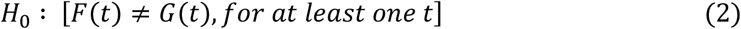

In order to compute the two-sided Kolmogorov–Smirnov alternative statistic for the above hypothesis, we first need to obtain the empirical cumulative distribution functions for both *X* and *Y* samples, such that for every real number *t*:

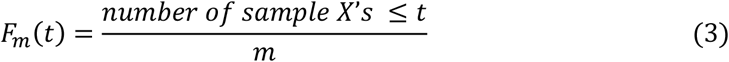

and

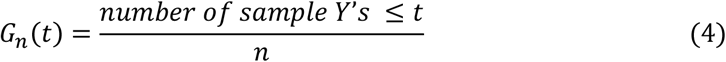

So, the two-sided two-sample Kolmogorov–Smirnov statistic is defined as:

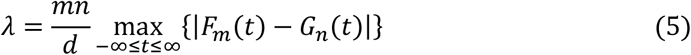

where *d* is the greatest common divisor of *m* and *n*.

Now, let’s consider the two images *m*_1_ and *m*_2_ with pixels *P*_1_ × *Q*_1_ and *P*_2_ × *Q*_2_ respectively, and suppose that *P*_1_ = *P*_2_ = *Q*_1_ = *Q*_2_ = 100 pixels, which means that we divide each axis for each image to 100 pixels. Then, in each pixel we count how many points/ molecules exist which gives the distribution for the points located in the *p*^th^ pixel. Thus, the empirical cumulative distribution for the points located in the *p*^th^ pixel for the two images can be defined as *F* = {*F_p_*, *p* = 0,…, 100} and *G* = {*G_p_*, *p* = 0,…, 100} respectively. Consequently, the distance of one distribution from the other is defined as:

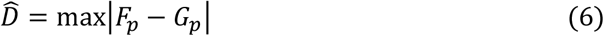

Under the assumption that the theoretical distributions for the two populations are the same, Kolmogorov and Smirnov [4] proved that the probability that the observed distance, 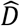 is greater than *D* is:

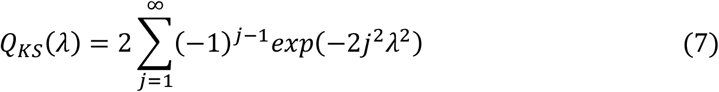

where λ is the Kolmogorov–Smirnov statistic:

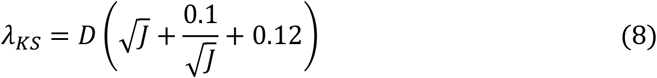

and

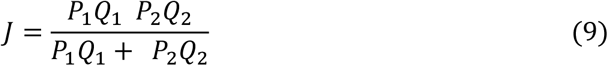

Thus, the test statistic *λ_KS_* defined in (8) can be used as a dissimilarity measure where the greater the distance between distributions, the bigger the value of *λ_KS_*. As a result, if we have a third image *m*_3_, we could say that both *m*_1_ and *m*_3_ are more similar to each other than *m*_2_ and *m*_3_ provided that *λ_KS_*(*m*_1_, *m*_3_) < *λ_KS_*(*m*_2_, *m*_3_).

More generally, assuming we have *k* groups of images/data sets, with population distributions *F*_1_, *F*_2_,…, *F_k_*, we may assign the image *m* to the group in which the mean of the dissimilarity values based on *F_i_* is smallest such that:

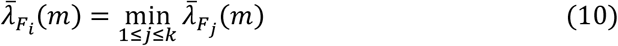

where *i* ≠ *j*, 1 ≤ *i* ≤ *k* and 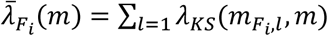.

### Significance testing

The p-values were generated via parametric, two-sided z-test using Graphpad Prism.

## Supplementary Information

**Supplementary Figure 1:**
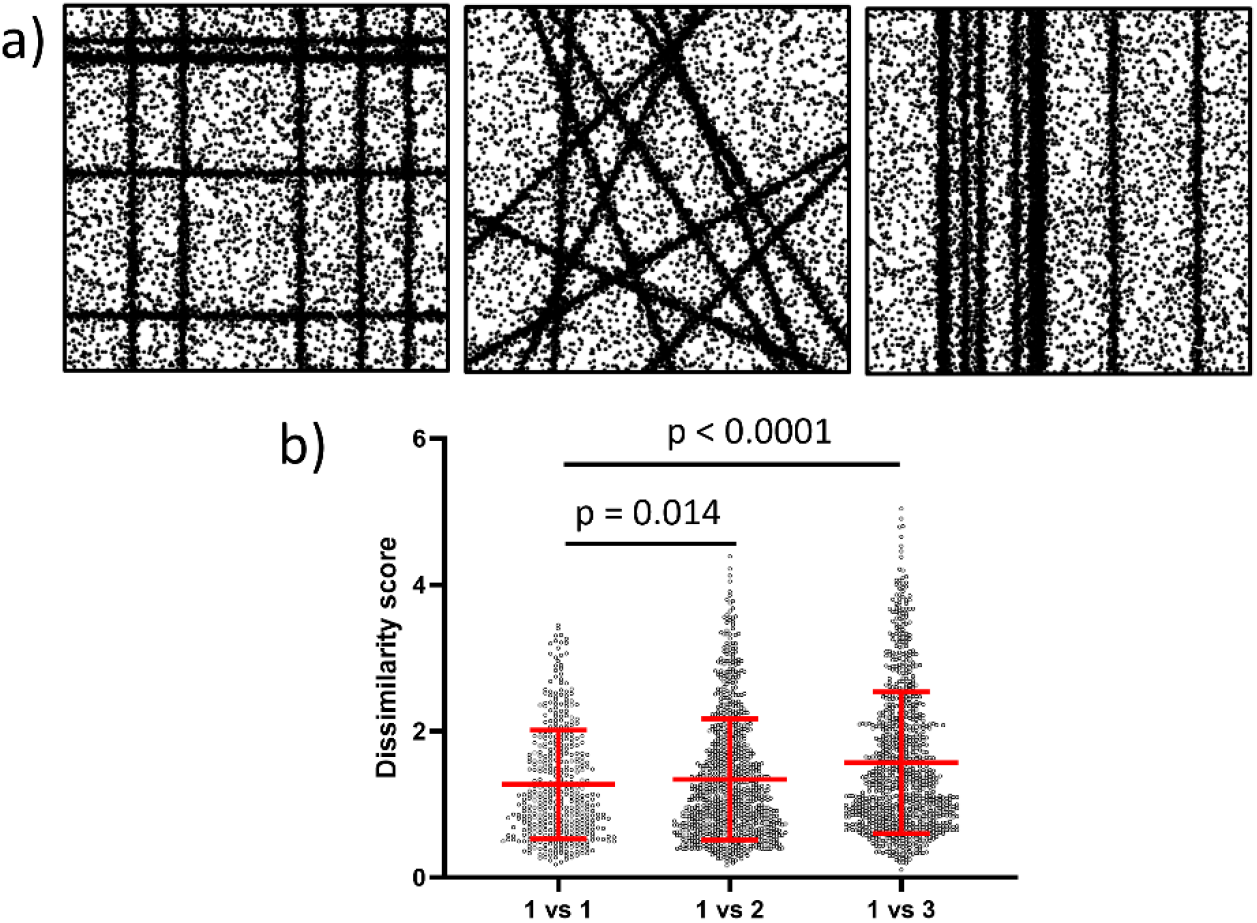
Similarity scores between point-clouds derived from simulated fibrous structures. a) Representative examples of 3000 x 3000 nm ROIs containing point-clouds derived from fibres placed on an orthogonal axis (Condition 1, left), randomly arranged linear fibres (Condition 2, centre) and parallel fibres (Condition 3, right). b) Dissimilarity scores between Condition 1 and conditions 2 and 3 show that the algorithm can discriminate between these different point-cloud distributions

**Supplementary Figure 2:**
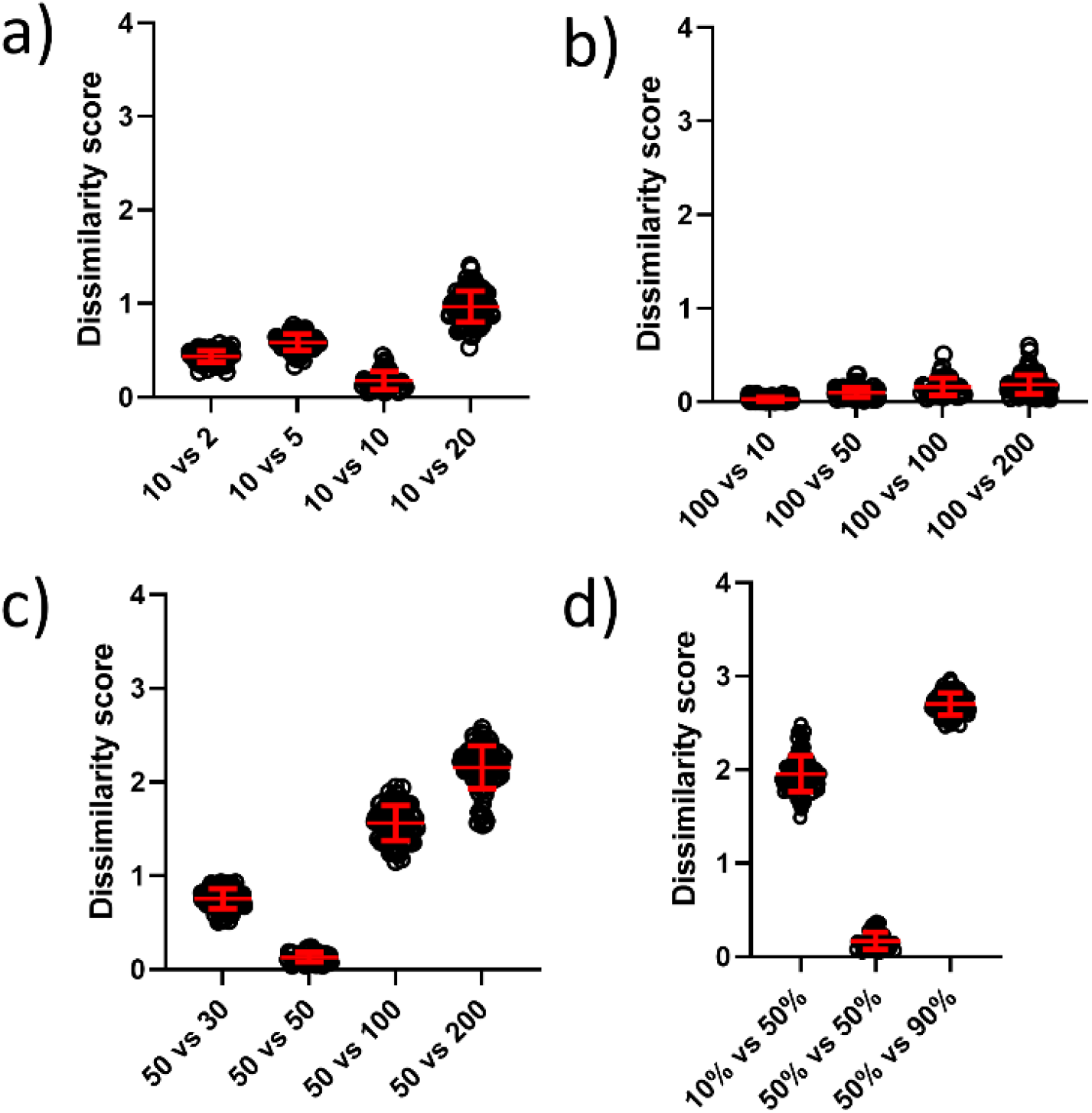
Demonstration of the method invariant to the total number of points on simulated data. Bee-swarm plots of λ-values for the default condition versus a) varying numbers of clusters per ROI. b) varying values of localisations per cluster. c) varying cluster sizes (nm). e) varying values for the percentage of un-clustered molecules. Bars show mean and S.D.

