## Supplementary material for "Measuring the similarity of SMLM-derived point-clouds": Software: mean_lambda.R

```
### pts.pix R 
### mean_lambda R 


#######################################################################################################################################################
#														
# TITLE:			Kolmogorov-Smirnov dissimilarity statistic for image comparison				
# AUTHOR:			Mohammed Baragilly 										
# DATE CREATED:			03/03/2022							   				
# DATE MODIFIED:		08/07/2022								   						
# PURPOSE: 			Function for calculating the mean of dissimilarities between different datasets from same or different conditions  	
#									 																														
#######################################################################################################################################################


pts.pix <- function(data, no.pix){
  sq <- seq(0, 3000, length.out = (no.pix+1))
  pix.counts <- NULL
  for(ii in 1:no.pix){
    for(jj in 1:no.pix){
      pix.counts <- c(pix.counts, length(which(sq[ii] <= data[,1] & sq[ii+1] >= data[,1] & sq[jj] <= data[,2] & sq[jj+1] >= data[,2])))
    }
  }
  pix.counts
}


mean_lambda <- function(folde1, folder2){
  if(folder1 == folder2){
    num.pxs=100	#number of pixels for each axis
    num_files <- length(list.files(folder1))
    file_i<-list.files(folder1)
    lambda <- NULL
    for(id in 1:(num_files-1)){
      for(jd in (id+1):num_files){
        sim1<-read.csv(paste(file=folder1,"/",file_i[id],sep=""),sep=",")
        xdata <- as.matrix(sim1[,c(1,2)])
        d1 <- pts.pix(xdata, num.pxs)
        nr <- num.pxs
        nc <- num.pxs
        J1 <- nr * nc
        sim2<-read.csv(paste(file=folder1,"/",file_i[jd],sep=""),sep=",")
        xdata2 <- as.matrix(sim2[,c(1,2)])
        d2 <- pts.pix(xdata2, num.pxs)
        nr <- num.pxs
        nc <- num.pxs
        J2 <- nr * nc
        h1 <- f1 <- h2 <- f2 <- rep(0, (num.pxs^2+1))
        for(i in 0:num.pxs^2) {
          h1[i + 1] <- length(d1[d1 == i])/length(d1)
          h2[i + 1] <- length(d2[d2 == i])/length(d2)
        }
        for(i in 2:(num.pxs^2+1)) {
          f1[i] <- f1[i - 1] + h1[i]
          f2[i] <- f2[i - 1] + h2[i]
        }
        f1[(num.pxs^2+1)] <- f2[(num.pxs^2+1)] <- 1
        mf <- max(abs(f1 - f2))
        J <- (J1 * J2)/(J1 + J2)
        lambdai <- mf * (sqrt(J) + 0.11/sqrt(J) + 0.12)
        lambda <- c(lambda, lambdai)
      }
    }
    out = list()
    out$lambda = lambda
    out$mean = mean(lambda)
    return(out)
  }
  if(folder1 != folder2){
    num.pxs=100	#number of pixels for each axis
    num_files1 <- length(list.files(folder1))
    num_files2 <- length(list.files(folder2))
    file1_i<-list.files(folder1)
    file2_i<-list.files(folder2)
    lambda <- NULL
    for(id in 1:num_files1){
      for(jd in 1:num_files2){
        sim1<-read.csv(paste(file=folder1,"/",file1_i[id],sep=""),sep=",")
        xdata <- as.matrix(sim1[,c(1,2)])
        d1 <- pts.pix(xdata, num.pxs)
        nr <- num.pxs
        nc <- num.pxs
        J1 <- nr * nc
        sim2<-read.csv(paste(file=folder2,"/",file2_i[jd],sep=""),sep=",")
        xdata2 <- as.matrix(sim2[,c(1,2)])
        d2 <- pts.pix(xdata2, num.pxs)
        nr <- num.pxs
        nc <- num.pxs
        J2 <- nr * nc
        h1 <- f1 <- h2 <- f2 <- rep(0, (num.pxs^2+1))
        for(i in 0:num.pxs^2) {
          h1[i + 1] <- length(d1[d1 == i])/length(d1)
          h2[i + 1] <- length(d2[d2 == i])/length(d2)
        }
        for(i in 2:(num.pxs^2+1)) {
          f1[i] <- f1[i - 1] + h1[i]
          f2[i] <- f2[i - 1] + h2[i]
        }
        f1[(num.pxs^2+1)] <- f2[(num.pxs^2+1)] <- 1
        mf <- max(abs(f1 - f2))
        J <- (J1 * J2)/(J1 + J2)
        lambdai <- mf * (sqrt(J) + 0.11/sqrt(J) + 0.12)
        lambda <- c(lambda, lambdai)
      }
    }
    out = list()
    out$lambda = lambda
    out$mean = mean(lambda)
    return(out)
  }
}


folder1=" C:/user/condition1 "
folder2=" C:/user/condition2 "


mean_lambda(folder1, folder2)
```
