## Supplementary material for "Measuring the similarity of SMLM-derived point-clouds": Software: User guide.docx

\### pts.pix R

\### mean_lambda R and mean_lambda_data_thinning R

TITLE:	Kolmogorov-Smirnov dissimilarity statistic for image comparison

AUTHOR:	Mohammed Baragilly

DATE CREATED:	03/03/2022

DATE MODIFIED:	08/07/2022

PURPOSE:	Function for calculating the mean of dissimilarities between different datasets from same or different conditions

***pts.pix***: it is a function to dividing the bivariate data matrix into specific number of pixels and then count how many points/ molecules exist in each pixel to get the distribution for the points located in divided pixels. It is used in the main function ***mean_lambda*** to get the related empirical cumulative distribution.

***mean_lambda***: it is a function to calculate the mean of dissimilarities between different datasets from same or different conditions. This is based on Kolmogorov-Smirnov test statistic by finding the empirical cumulative distribution functions for two samples.

***mean_lambda_data_thinning***: is the total-points-invariant version

**User guide:**

In order to use the function ***mean_lambda or mean_lambda_data_thinning***, we have to identify the location of the folder contains the data files to be compared.

First of all, we need to recall the two functions ***pts.pix*** and ***mean_lambda*** by selecting all their lines and then right click and choose Run line or selection.

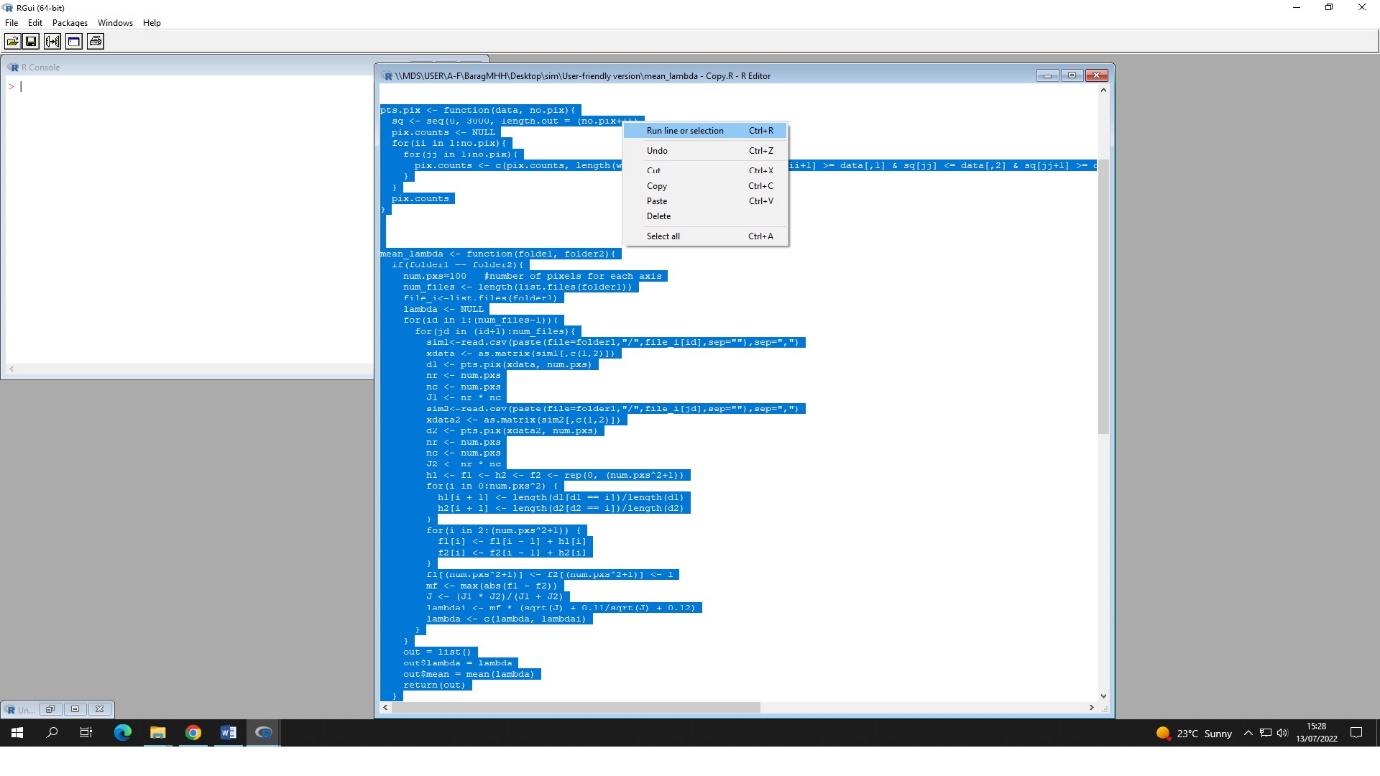

There are two options to be considered before using the function:

(A) We compare different datasets from same condition (i.e. all the files are located in one folder). So, in this case we identify the path of folder1 to be exactly same as the path of folder2. For example:

*folder1=" C:/user/condition1 "*

*folder2=" C:/user/condition1 "*

(B) We compare different datasets from different conditions (i.e. the data files for the first condition are located in some folder and the data files for the second condition are located in different folder). So, in this case we identify the path of each folder. For example:

*folder1=" C:/user/condition1 "*

*folder2=" C:/user/condition2 "*

Once we added the location/path of folder1 and folder2, we can now run the function ***mean_lambda*** or ***mean_lambda_data_thinning*** as following:

*mean_lambda(folder1, folder2)*

*mean_lambda_data_thinning(folder1, folder2)*

Please note that data files should include x and y values in the first two columns.
